# Evaluation of Tumor-colonizing *Salmonella* Strains using the Chick Chorioallantoic Membrane Model

**DOI:** 10.1101/2024.10.27.620524

**Authors:** Khin K. Z. Mon, Linda J. Kenney

## Abstract

The chick embryo chorioallantoic membrane (CAM) tumor model is a valuable preclinical model for studying the tumor colonizing process of *Salmonella enterica* serovar Typhimurium. It offers advantages such as cost-effectiveness, rapid turnaround, reduced engraftment issues, and ease of observation. In this study, we explored and validated the applicability of the partially immune-deficient CAM tumor model. Herein, we demonstrate that *Salmonella* preferentially colonizes tumors and directly causes tumor cell death. Bacterial migration, tumor colonization, and intra-tumor distribution did not require flagellar-mediated motility. The vast majority of *Salmonella* that colonized the CAM tumor were extracellular. Thus, tumor invasion was independent of both SPI1– and SPI2-encoded type three secretion systems. Surprisingly, the extracellular residence of *Salmonella* on CAM tumors did not require biofilm formation. We evaluated our wild-type parental strain compared to the attenuated clinical strain VNP20009 and discovered a reduced tumor colonization capability of VNP20009. The inability to effectively colonize CAM tumors potentially explains the reduced anti-tumor efficacy of VNP20009. Our work establishes the xenograft CAM model as an informative and predictive screening platform for studying tumor-colonizing *Salmonella*.

**IMPORTANCE:** Cancer has a major impact on society, as it poses a significant health burden to human populations worldwide. *Salmonella* Typhimurium has demonstrated promise in cancer treatment by exerting direct tumoricidal effects and enhancing host-mediated anti-tumor immunity in xenograft mouse studies. A general understanding of its pathogenesis and the relative ease of genetic manipulation support the development of attenuated strains for therapeutic use. Alternative *in ovo* models such as the CAM tumor model present a suitable screening platform to accelerate the development of therapeutic strains. It allows for rapid evaluation of *Salmonella* strains to assess their efficacy and potential as oncolytic agents. The present study establishes that the *in ovo* tumor model can be utilized as a preclinical tool for evaluating oncolytic *Salmonella*, bridging the gap between *in vitro* and *in vivo* screening.

## INTRODUCTION

In the field of bacterial-mediated cancer therapy, *Salmonella enterica* serovar Typhimurium (STm) has emerged as an ideal therapeutic candidate. This is in part due to its intrinsic tumor targeting ability (1), its facultative anaerobic properties, which allow colonization in both hypoxic and oxygenated regions of heterogeneous tumor microenvironments (2), its invasive capability (3), and its direct or immune-mediated anti-tumor effects (3–6). STm infection mainly causes self-limiting gastrointestinal disease in healthy humans but can sometimes manifest as extraintestinal systemic infections in immune-compromised individuals (7). To promote its safe application in cancer patients, bacterial pathogenic properties need to be eliminated or attenuated without compromising the tumor colonizing capability, and thus subsequent anti-tumor potency. Therefore, tuning the balance between virulence attenuation and tumor-colonization capability is an important criterion in the design of suitable STm strains.

The chick embryo chorioallantoic membrane (CAM) has long been used as a robust model for cancer and cardiovascular research, as well as for preclinical evaluation of cancer drug therapies (8–10). The CAM model is an immunodeficient model that is receptive to xenografting of various mammalian tumor cells without inducing a host immune response (11). The well-vascularized embryonic tissue of the CAM supports and preserves heterogeneous tumor growth with a much shorter experimental timeline (typically 3-4 days) for tumor development compared to mouse models, which requires weeks (12). Moreover, the CAM model also allows real-time visualization of tumor growth (13) as well as the dynamics of bacterial infection via fluorescence stereomicroscopy (14). These attributes indicate that the CAM tumor model could be well suited for preliminary evaluation of *Salmonella*-based therapies for cancer treatment.

The engineered STm strain VNP20009 is attenuated by purine auxotrophy (a deletion in *purI*) and lipid A modification (a deletion in *msbB*) (15). It demonstrated an excellent safety profile with observable tumor growth retardation in a variety of pre-clinical xenograft mouse models (16–19). Despite the high success rate in the mouse studies, no anti-tumor effect was observed in humans. The intravenous infusion of the VNP20009 strain to melanoma patients in a clinical trial failed and was attributed to a low bacterial tumor colonization and rapid bacterial clearance by the host immune response (20). The failure of the VNP20009 strain to colonize tumors highlighted the xenograft mouse model as a poor predictor of therapeutic success in humans. Hence, there is a need to explore alternative models to overcome some of the translational limitations associated with traditional mouse models. We hypothesized that the CAM tumor model might serve as a preliminary screening platform to study the tumor-bacterial interactions to evaluate the efficacy of tumor-targeted *Salmonella* strains for therapeutic purposes.

In the current work, we developed the *in ovo* CAM tumor model and compared it with an *in vitro* three-dimensional (3D) tumor spheroid model to examine the relevance of the screening data obtained (21). While 3D tumor spheroids can mimic *in vivo* solid tumor features such as structural organization, cellular assembly and nutrient gradients (22), they cannot fully simulate the heterogenous tumor microenvironment established within a living host. Therefore, the CAM tumor model serves as a better pre-clinical model, bridging an informational gap between *in vitro* cell-based models and *in vivo* animal-based systems for screening oncolytic *Salmonella.* With the CAM tumor model, we demonstrated that STm pathogenic traits such as flagella, virulence factors and biofilm formation were not essential for tumor colonization. We further discovered that reduced tumor colonization by VNP20009 was attributed to its growth sensitivity within the tumor microenvironment, which subsequently impacted its anti-tumor effectiveness.

## RESULTS

### Validation of the CAM model for studying tumor-colonizing *Salmonella*

A general overview of the CAM tumor STm infection model is depicted in Figure 1. We first determined the infectious dose of STm by infecting tumor bearing chick embryos with different doses of bacteria: approximately 500 colony-forming units (CFU), 50 CFU, or 5 CFU. Chick embryo mortality was monitored daily post-bacterial infection with PBS as the control group. The highest inoculum of ∼500 CFU caused the greatest mortality (38% survival) at one day post-infection (DPI) compared to groups that received lower dosages of ∼50 CFU (66% survival) or ∼5 CFU (77% survival) (Supplementary Figure 1). To minimize the mortality of highly susceptible tumor-bearing chick embryos as well as to ensure effective tumor colonization, we selected the mid-range of ∼50 CFU as the optimal infectious dose.

**Figure 1:**
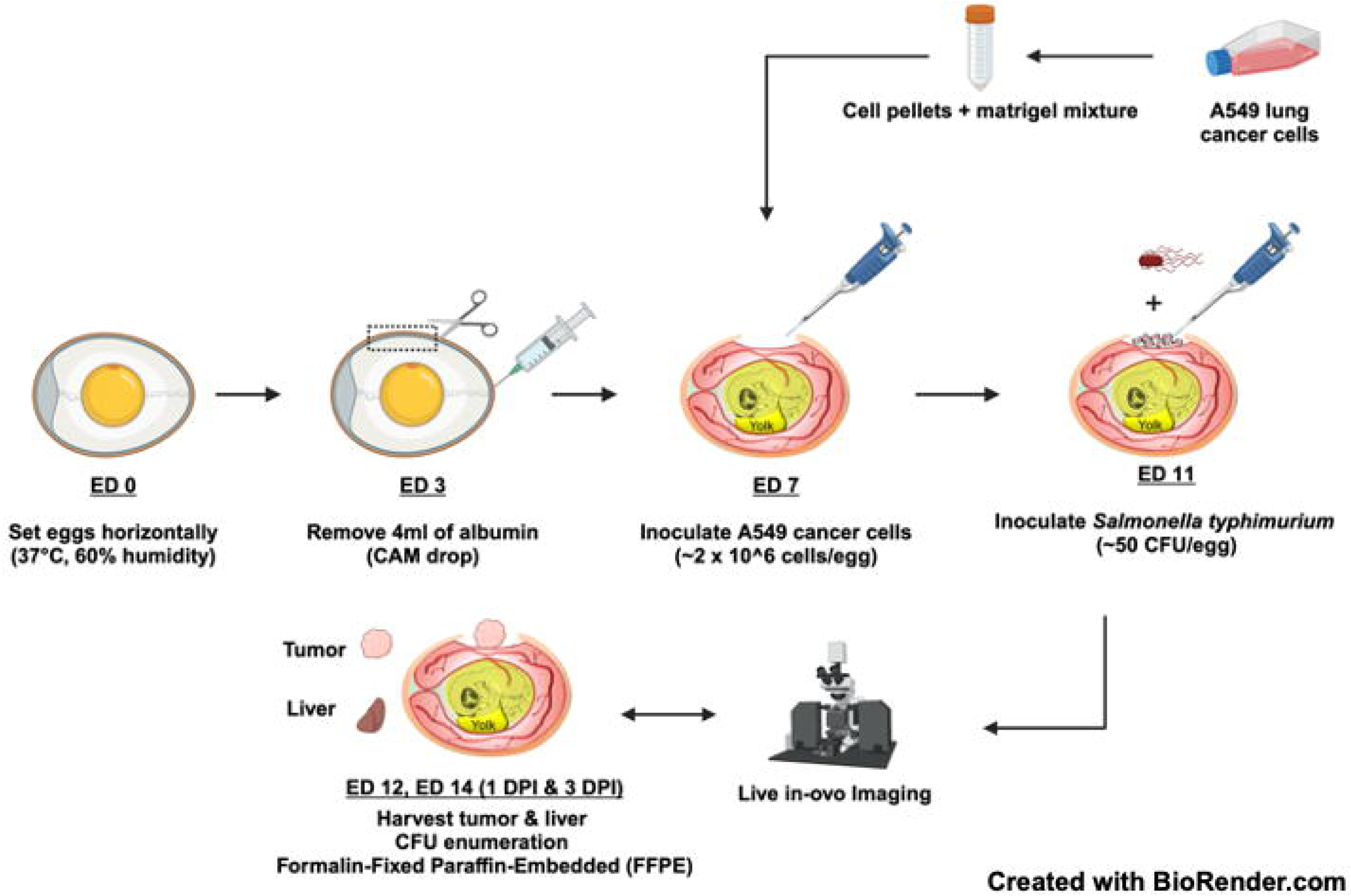
Experimental scheme of the CAM tumor, STm infection model.

The important features of tumor-targeting STm are the higher tumor specificity in colonization compared to organ colonization, and the ability to penetrate tumor tissue (23, 24). These criteria were tested first in the CAM tumor model. At 1 DPI, WT STm accumulated preferentially in tumors over livers of chick embryos at a ratio of 1000:1. The mean CFU/g in tumors was 1.9 x 10^8^ CFU/g compared to 1.9 x 10^5^ CFU/g in livers (Figure 2A, note log scale). Next, we examined the intra-tumoral distribution of STm. Infected whole tumors were harvested, embedded in paraffin and sectioned at different tissue depths (top, middle and bottom). The tumor sections were stained with antibodies against bacterial lipopolysaccharide (green), nuclei (blue) and E-cadherin (magenta) and examined by confocal microscopy. STm was observed at all tumor tissue depths as evident by lipopolysaccharide staining (LPS = green, second panel Figure 2B and Supplementary Figure 2). Specifically, bacteria were mostly present in the metastatic tumor cell regions, where decreased E-cadherin staining indicated a loss of cell-to-cell adhesion junctions.

**Figure 2:**
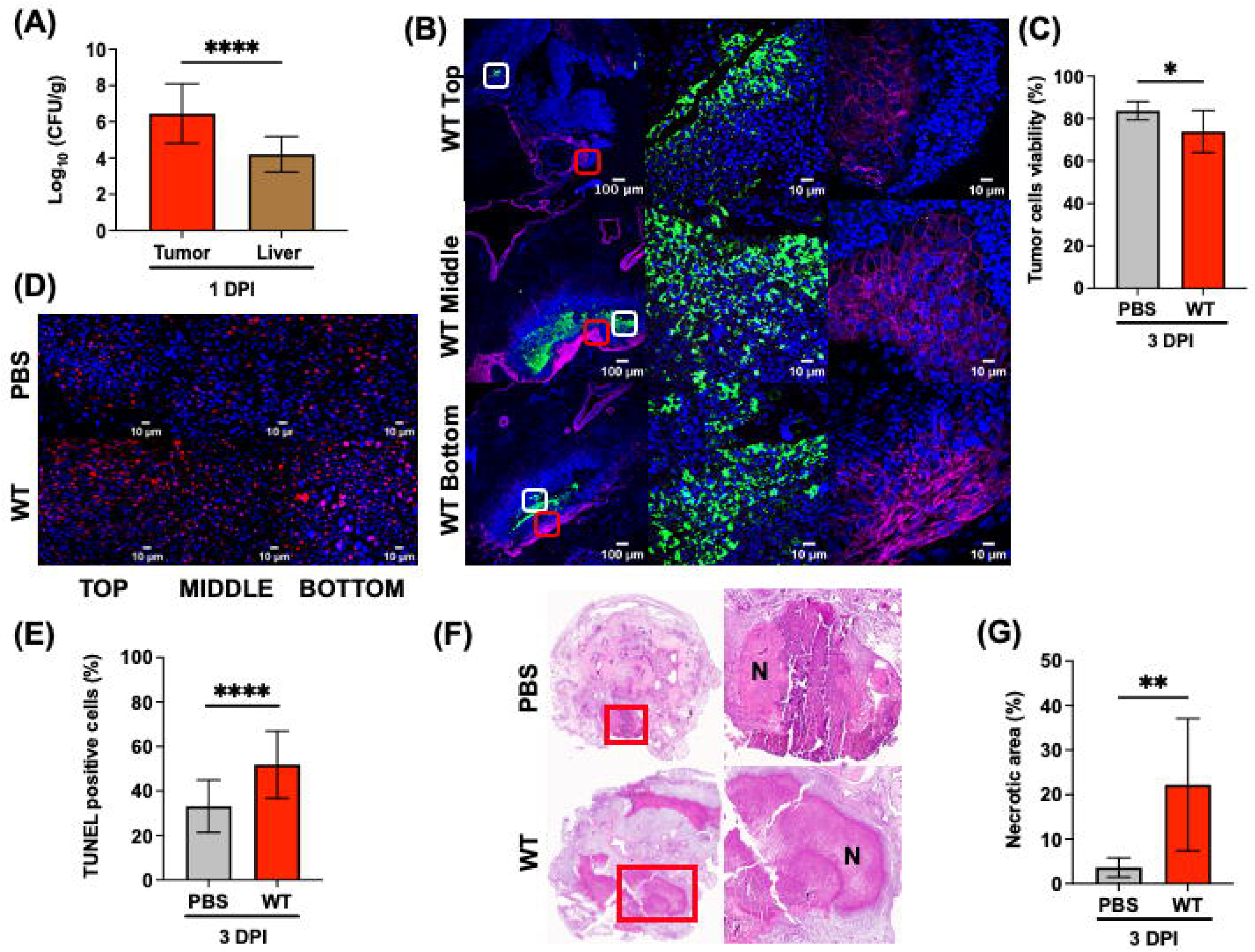
S**T**m **promotes tumor cell death in the CAM model.** (A-G) Chick embryos with pre-established A549 tumors were directly inoculated with *Salmonella* WT or PBS on ED 11. Tumors and livers were harvested at 1 and 3 DPI. (A) *Salmonella* WT CFU recovered per gram of tissue from tumors and livers at 1 DPI (tumor to liver ratio 1000:1). (B) Representative images of *Salmonella* WT LPS (green), E-cadherin (magenta) and Hoechst (blue) staining detected in tumor sections from top, middle and bottom of whole tumors harvested at 1 DPI. Scale bar = 100 µm (10x objective) and 10 µm (100x objective). The middle panel is the zoomed image of the white box and the right panel is the zoomed image of the red box at 100x objective. (C) A trypan blue exclusion assay was performed on *ex vivo* dissociated tumor cell suspensions, indicating a loss of tumor cell viability in the group infected with WT *Salmonella* at 3 DPI. (D) Representative images of TUNEL-stained (red) and Hoechst (blue) tumor sections from 3 DPI (top, middle, bottom). Scale bar = 10 μm, 100x objective. (E) Quantitative comparison of the percentage of TUNEL-positive nuclei between WT-infected tumors (52%) and PBS-treated tumors (33%) at 3 DPI. (F) Representative images of H&E tumor sections. Zoomed images of the red box of the necrotic region at 3 DPI. N: necrotic areas. (G) The percentage of the necrotic area as quantified by an external pathologist from H and E-stained tumor sections infected with WT (22%) and PBS (4%) at 3 DPI. The results represent the mean ± SD. For all, the statistical significance was by unpaired t-test, not significant (ns). P-value * ≤ 0.05, ** ≤ 0.01, *** ≤ 0.001, **** ≤ 0.0001.

### STm colonization drove direct tumor apoptosis in the CAM model

We next tested the hypothesis that STm infection directly promotes tumor cell death. A trypan blue exclusion assay was performed on dissociated tumor cells to test *ex vivo* tumor cell viability. At 3 DPI, STm infection significantly reduced the percentage of tumor cell viability (74%) compared to PBS treated tumors (84%) (Figure 2C). Because the presence of STm was detected throughout the entire tumor (Figure 2B), it raised the question of whether its localization directly induced tumor cell apoptosis. We utilized the terminal deoxynucleotidyl transferase-mediated deoxyuridine triphosphate nick end-labeling (TUNEL) assay to detect *Salmonella*-induced DNA fragmentation in tumor cells. A marked accumulation of TUNEL-positive cells (red signals) was observed in STm-infected tumors relative to PBS-treated tumor sections at the top, middle and bottom levels (Figure 2D). Immunofluorescence images of TUNEL-stained tumor sections were further quantified using Fiji software (25). The percentage of TUNEL-positive nuclei in STm infected tumor tissue sections was significantly higher than PBS tumors (52% vs 33%, respectively), indicating that tumor cells underwent increased apoptosis following bacterial infection (Figure 2E). Lastly, the histological changes in Hematoxylin and Eosin (H&E)-stained tumor sections were analyzed. A more extensive necrotic area in *Salmonella*-infected tumors (22%) was detected compared to the PBS control group (4%) (Figure 2F, 2G).

### Bacterial motility was not required for tumor colonization or intra-tumoral distribution

The prevailing hypothesis regarding a role for flagella in tumor colonization is that motility is essential for bacteria to effectively target, colonize, and disperse within the tumor environment (26–28). To test this hypothesis, WT or a Δ*flgK* null strain were directly inoculated onto CAM tumors. The *flgK* gene encodes a flagellar hook-associated protein; its deletion renders *Salmonella* non-motile (29). Live *in ovo* images of the bacterial colonization pattern, as well as CFU enumeration from tumors was comparable between the WT and the Δ*flgK* null strain (Figure 3A, B). We reasoned that the role of flagella might be dispensable when bacteria were directly inoculated onto the tumor, rendering colonization a passive process. Therefore, we also tested a distant inoculation route by depositing bacteria a short distance away from the tumor (∼0.5 cm) on filter papers to test whether *Salmonella* required flagellar-mediated motility for tumor colonization. The Δ*flgK* strain showed no defect in CAM tumor colonization, and colonized tumors to the same extent as the WT (Figure 3A, C). The intra-tumoral distribution pattern of the non-motile Δ*flgK* strain was also comparable to that of the motile WT (Figure 2B), indicating that the Δ*flgK* strain was able to disperse and penetrate all depths of the tumor (Figure 3D). These results provided strong evidence that STm migration, tumor colonization and intra-tumoral distribution were not dependent on bacterial motility in the CAM tumor model.

**Figure 3:**
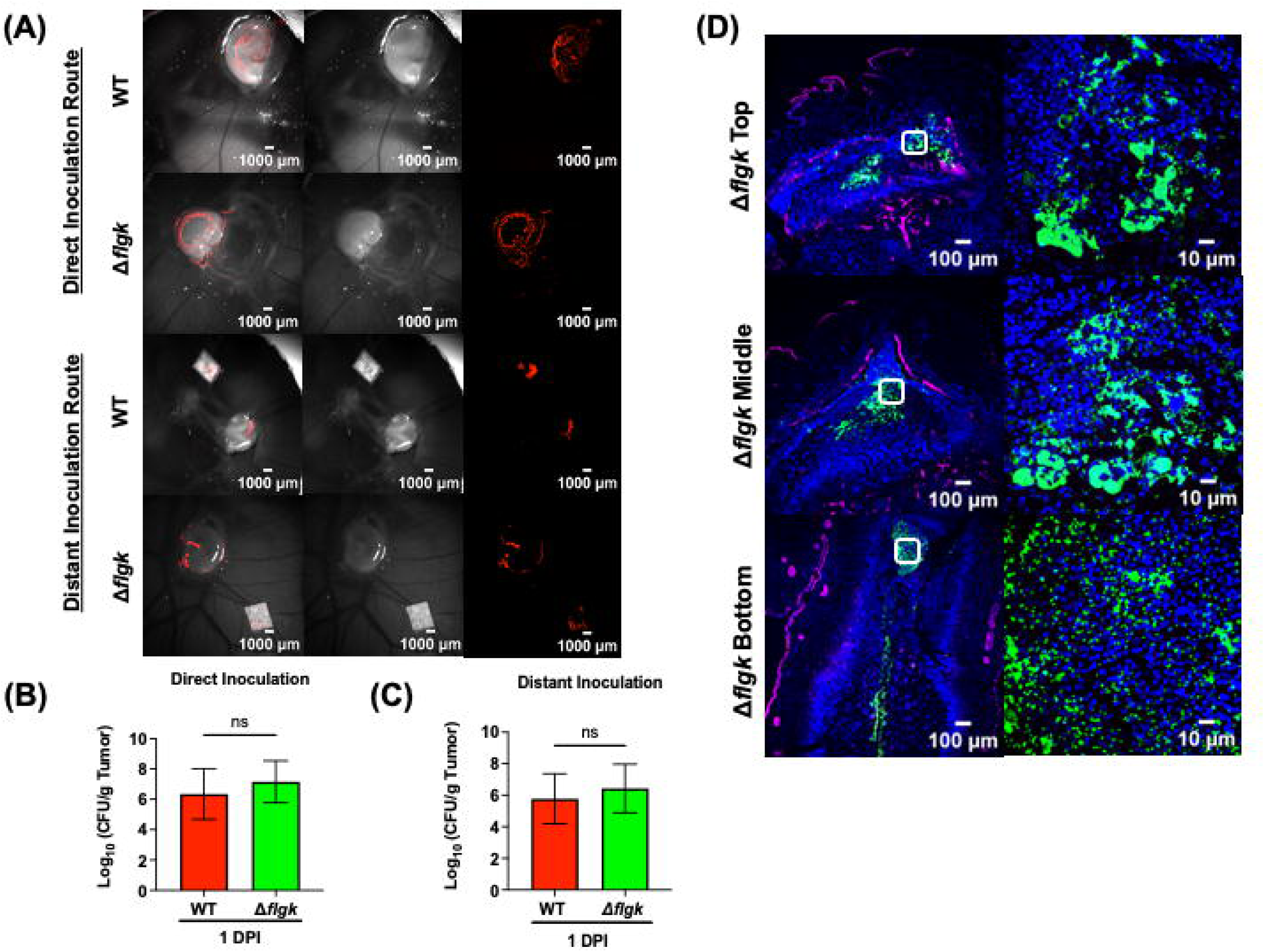
F**l**agellar **motility is not required for tumor colonization or intra-tumoral distribution.** (A) Representative live *in ovo* images of mCherry-tagged *Salmonella.* WT and Δ*flgK* colonizing A549 tumors on CAMs via either direct (bacteria inoculated directly on tumors) or distant inoculation (bacteria deposited on filter paper, ∼0.5 cm away from tumors) at 1 DPI. (B-C) Total *Salmonella* WT and Δ*flgK* CFU recovered per gram of tumor tissue at 1 DPI from direct and distant inoculation. The results represent the mean ± SD. The statistical significance was by unpaired t-test, not significant (ns). (D) Representative images of *Salmonella* Δ*flgK* LPS (green), E-cadherin (magenta) and Hoechst (blue) staining detected in tumor sections from top, middle and bottom levels of whole tumors harvested at 1 DPI. Scale bar = 100 µm (10x objective) and 10 µm (100x objective). The second column is the zoomed in details of the white box at 100x objective.

### 3D tumor spheroids do not recapitulate the CAM tumor model with respect to key virulence factors encoded on pathogenicity islands 1 and 2

In the host gastrointestinal tract, STm utilizes a Type III secretion system (T3SS1) encoded by *Salmonella* Pathogenicity Island-1 (SPI-1) to invade host intestinal epithelial cells and a SPI-2-encoded T3SS2 for intracellular growth and survival within host macrophages (30–33). We therefore investigated whether bacterial invasion or replication within tumors was also dependent on SPI-1 and SPI-2. To address this question, we first employed the *in vitro* 3D tumor spheroid model before assessing their roles *in vivo* in the CAM tumor model. The use of 3D magnetic cell culture technology produced structurally relevant tumor spheroids that mimicked some features of *in vivo* solid tumors. The spatial architecture with regard to oxygen and nutrient gradients, extracellular matrix, and cell-to-cell interactions is maintained (21, 22). Within tumor spheroids, intracellular invasion and replication rates of STm strains were examined using a gentamicin protection assay. The elimination of *hilD* (the SPI-1 master regulator) resulted in a significant invasion defect with a mean reduction of 1.4 log CFU/well compared to WT (Figure 4A), suggesting that SPI-1 played a significant role in early bacterial invasion of tumor cells. At 16 HPI, an intracellular replication rate of both Δ*hilD* (mean log CFU/well = 3.5) and the SPI-2 master regulator, Δ*ssrB* (mean log CFU/well = 4.5) were significantly reduced relative to WT (mean log CFU/well = 5.6) (Figure 4A). Consequently, the overall replication rate (16 HPI/ 2 HPI) was higher for WT (28-fold) compared to either Δ*hilD* and Δ*ssrB* strains (6-fold and 4-fold, respectively) (Figure 4B). We next examined the cytotoxic damage induced by bacterial infection of 3D tumor spheroids using a lactate dehydrogenase (LDH) assay 16 HPI. The amount of LDH release is directly correlated with cell cytotoxicity levels. Tumor spheroids infected with WT bacteria released higher amounts of LDH (53% of total maximum LDH release at 100%) compared to SPI-1 (22%) or SPI-2 (33%) deficient strains (Figure 4C). Whole-mount immunofluorescence staining of 3D tumor spheroids exhibited a specific cellular structural organization. Proliferating tumor cells were observed in the periphery, with an inner layer of quiescent cells and a necrotic core region. These were evident by distinct fluorescence intensity profiles of the cell proliferation marker Ki67, in which the intensity varied from high to low from the periphery to the core (Figure 4D). From representative images of infected 3D tumor spheroids at 16 HPI, both SPI-1– and SPI-2-deficient strains were observed at reduced levels compared to the WT (Figure 4D) consistent with the quantification of CFU (Figure 4A).

**Figure 4:**
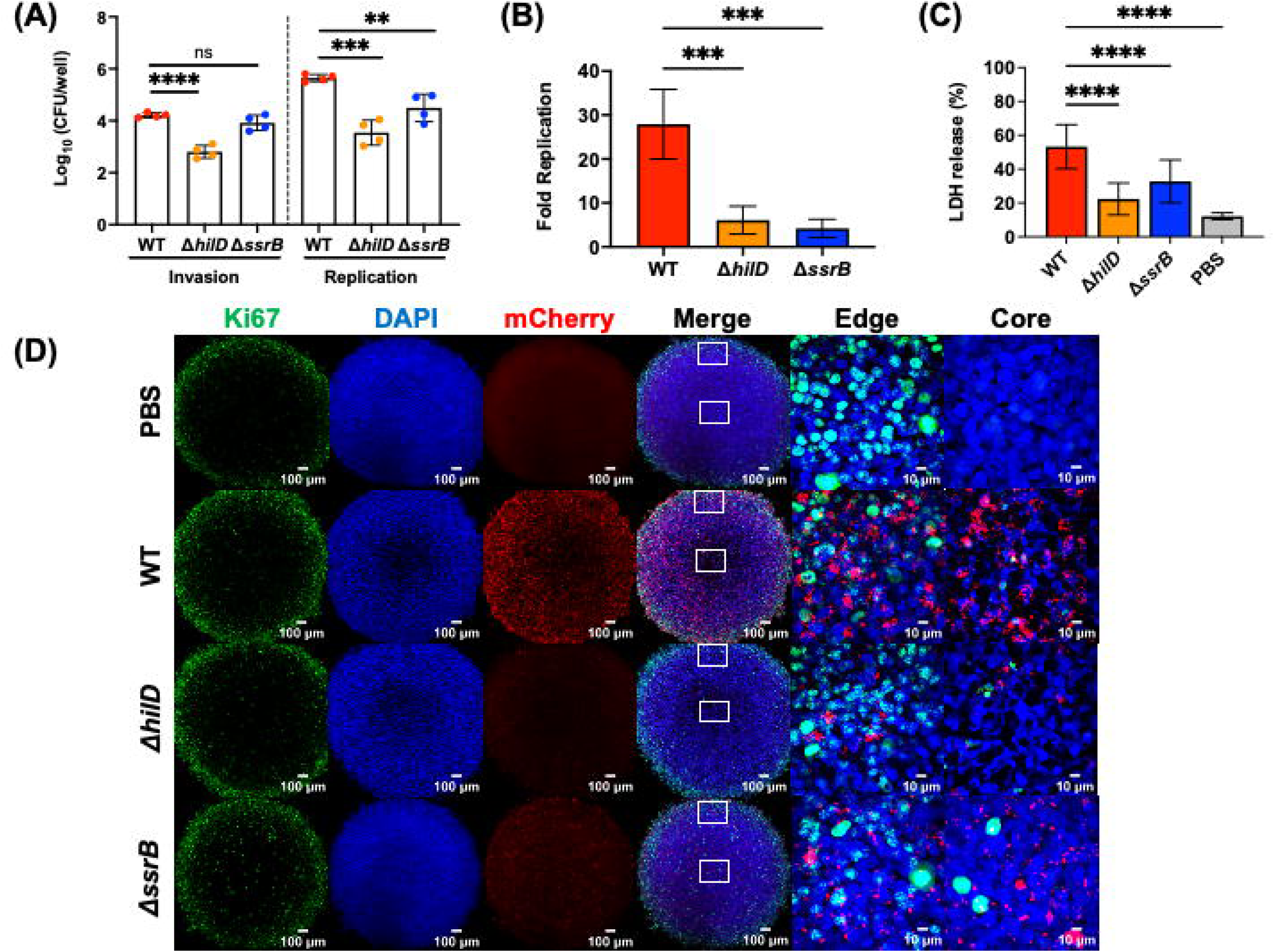
S**P**I**-1 and SPI-2 are required for intracellular replication of STm within 3D tumor spheroids.** (A) Intracellular bacterial enumeration of STm strains (WT, Δ*hilD and* Δ*ssrB)* in A549 tumor spheroids measured at invasion (mean log CFU/well at 2 HPI: WT = 4.2, Δ*hilD* = 2.8, Δ*ssrB* = 3.9) and replication (mean log CFU/well at 16 HPI: WT = 5.6, Δ*hilD* = 3.5, Δ*ssrB* = 4.5) stages of infection. Dots: biological replicates, each with a minimum of five technical replicates. The bars represent the mean +/− SD. (B) Fold replication (16 HPI/2 HPI) of each STm strain (WT = 28-fold, Δ*hilD* = 6-fold, Δ*ssrB* = 4-fold) within A549 tumor spheroids. The bars represent the means of four biological replicates, a minimum of five technical replicates each. Error bars +/− SD. (C) The amount of LDH released from A549 tumor spheroids after infection with STm (WT= 53%, Δ*hilD* = 22% and Δ*ssrB* = 33%) at 16 HPI. The bars represent the means of three biological replicates, a minimum of five technical replicates each. Error bars +/− SD. For all, the statistical significance was determined by one way analysis of variance (ANOVA) with Sidak’s multiple comparison test. Not significant (ns), P-value *≤ 0.05, **≤ 0.01, ***≤ 0.001, ****≤ 0.0001. (D) Representative images of immunofluorescence-labeling of 3D tumor spheroids with Ki67 AlexaFluor 488 (green), Hoeschst (blue) and mCherry-tagged bacterial strains (red) at both 10x and 100x objectives (edge and core of spheroids). Maximum intensity projections of Z-stack images were taken with an Olympus Super Resolution Spinning Disk SpinSR-10 microscope. Scale bar = 100 μm (10x) and 10 μm (100x).

We then investigated whether tumor colonization and intracellular invasion or replication required SPI-1 or SPI-2 in the CAM tumor model. Using an ex vivo gentamicin assay, we quantified total tumor-associated bacteria (i.e. both extra– and intra-cellular bacteria) recovered from the CAM tumors. Both SPI-1 (Δ*hilD)* and SPI-2 (Δ*ssrB*)-deficient strains were found to colonize (extracellular) and invade (intracellular) tumor cells to the same extent as WT at 1 DPI (Figure 5A, B). In the mouse tumor model, *Salmonella* is found almost exclusively extracellularly, forming biofilms on tumors (34). We therefore assessed whether *Salmonella* adapted a similar residence lifestyle in our CAM tumor model. Based on the percentage of the total tumor-associated bacterial population, approximately 99% of bacteria were extracellular, with ∼1% of the bacteria found inside the CAM tumor cells (WT, Δ*hilD* and Δ*ssrB*) (Figure 5C). Hence, this finding corroborated that the loss of SPI-1 and SPI-2 genes did not impair the tumor-colonizing ability of STm (Figure 5B), because STm were predominantly extracellular on the CAM tumors.

**Figure 5:**
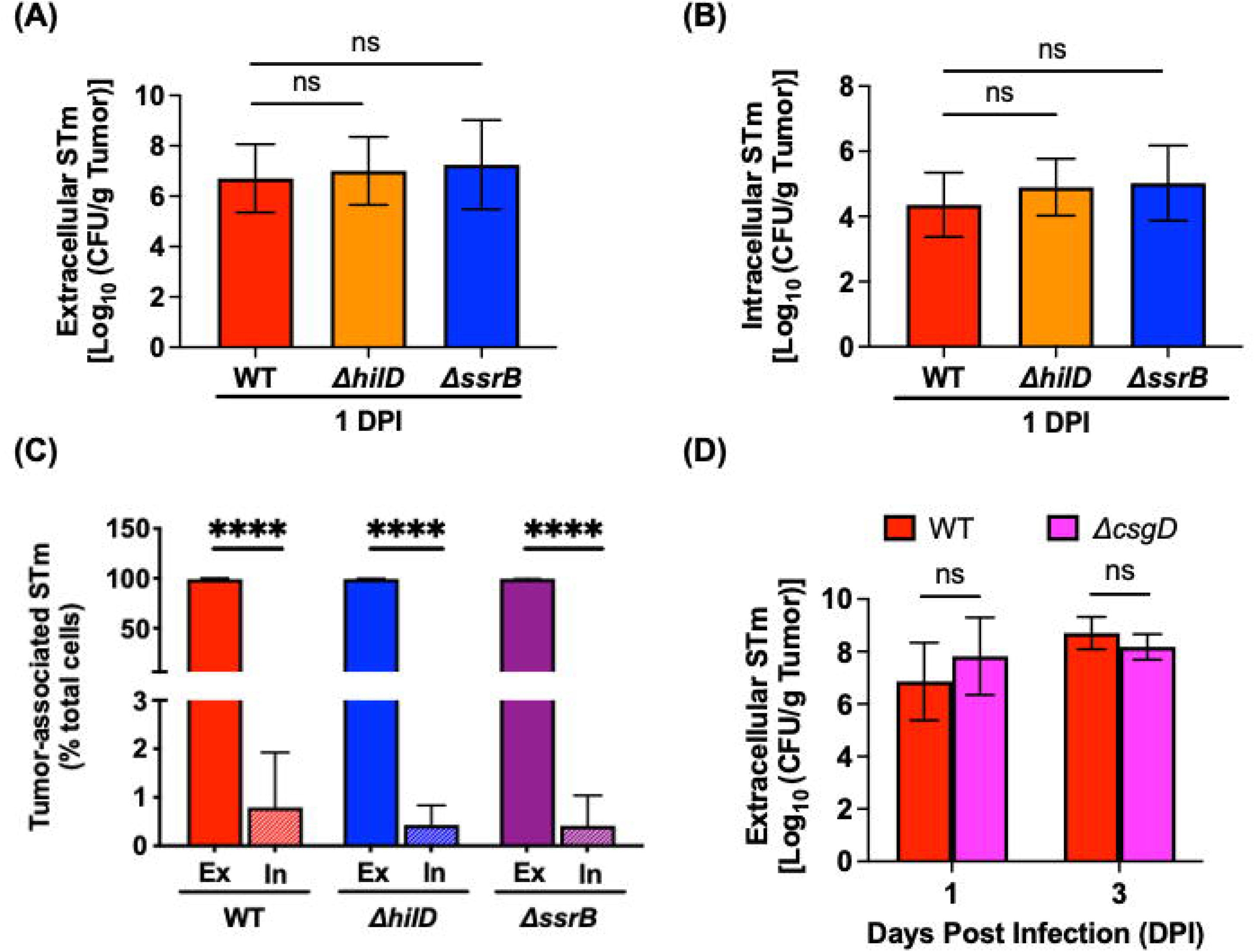
I**n the CAM tumor model, STm resides in an extracellular, non-biofilm state.** (A) Extracellular CFU recovered after infection with WT *Salmonella*, Δ*hilD* and Δ*ssrB* per gram of tumor at 1 DPI. Statistical significance by one-way ANOVA with Sidak’s multiple comparison test. (B) Intracellular WT *Salmonella*, Δ*hilD* and Δ*ssrB* CFU recovered per gram of tumor at 1 DPI. Statistical significance by one-way ANOVA with Sidak’s multiple comparison test. (C) Percentage of total tumor-associated STm (WT, Δ*hilD,* Δ*ssrB)* recovered before and after an ex vivo gentamicin assay from infected CAM tumors at 1 DPI. EX: extracellular, IN: intracellular. Statistical significance was by one-way ANOVA with Sidak’s multiple comparison test. (D) Extracellular *Salmonella* WT and Δ*csgD* CFU recovered per gram of tumor at 1 and 3 DPI. The results represent the mean ± SD. The statistical significance was by unpaired t-test. For all, not significant (ns), P-value * ≤ 0.05, ** ≤ 0.01, *** ≤ 0.001, **** ≤ 0.0001.

To address the question of whether extracellular STm also form biofilms on CAM tumors, we infected tumors with *Salmonella* WT or a Δ*csgD* null strain; CsgD is the master biofilm regulator (35). The extracellular bacterial load recovered indicated that both strains successfully colonized the tumor to the same extent: a mean log CFU/g of ∼7 to 8, at 1 and 3 DPI (Figure 5D). Thus, in contrast to the mouse tumor model (34), the ability to form biofilms was not required for the extracellular accumulation of STm on CAM tumors.

### How does the clinical strain VNP2009 compare with WT *Salmonella*?

Despite promising results in xenograft mouse models (effective tumor colonization with tumor growth retardation) the clinical strain VNP20009 did not perform well (low bacterial tumor colonization) in a human clinical trial. It was therefore of interest to evaluate colonization by VNP20009 compared to its parental WT strain (15, 36) in both 3D tumor spheroid and CAM tumor models.

### The clinical strain VNP20009 replicates poorly within 3D tumor spheroids

We first investigated the performance of the clinical strain VNP20009 within 3D tumor spheroids. Compared to WT, the VNP20009 strain displayed a modest invasion defect with a mean reduction of 0.6 log CFU/well (Figure 6A, left columns). However, the replication defect at 16 HPI was striking, with an overall mean reduction of 1.6 log CFU/well relative to WT (Figure 6A, right columns). Replication of the WT was 28-fold, but VNP20009 only increased 3-fold (Figure 6B). These differences indicated a stark difference in the ability of the attenuated VNP20009 strain to grow within the tumor spheroid compared to the WT. The possibility that differential growth rates between strains contributed to tumor invasion was ruled out, as the growth rate of VNP20009 was not compromised under bacterial culture conditions (Supplementary Figure 3). Furthermore, bacterial inoculums were prepared under the same SPI-1 inducing condition for both strains. The percentage of bacteria that were SPI-1 positive (using a P*prgH*-mCherry reporter) for WT and VNP20009 were similar at 67% and 62%, respectively, of the total population (Supplementary Figure 4). We next compared the ability of VNP20009 to induce tumor cell cytotoxicity with the LDH assay after 16 HPI. WT-infected tumor spheroids released higher amounts of LDH (53%) than VNP20009 (36%) (Figure 6C).

**Figure 6:**
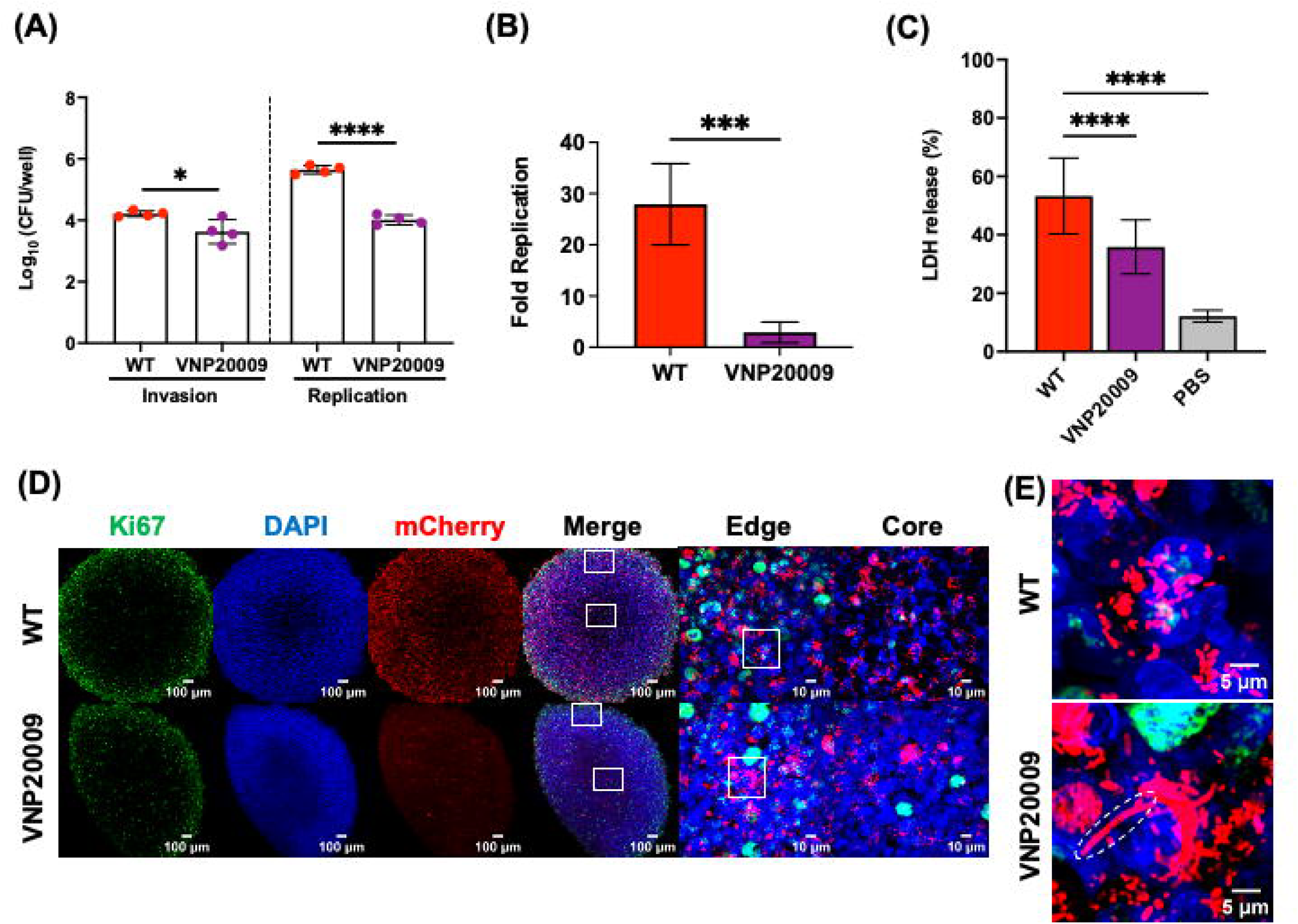
R**e**plication **of VNP20009 within 3D tumor spheroids is impaired.** (A) Intracellular bacterial enumeration from WT and VNP20009 infected tumor spheroids, measured at invasion (mean log CFU/well at 2 HPI: WT = 4.2, VNP20009 = 3.6) and replication (mean log CFU/well at 16 HPI: WT = 5.6, VNP20009 = 4) stages of infection. Dots: biological replicates, each with a minimum of five technical replicates. The bars represent the mean +/− SD. The statistical significance was determined by unpaired t-test. (B) Fold replication (16 HPI/2 HPI) of WT (28-fold) and VNP20009 (3-fold) strains within tumor spheroids. The bars represent the means of four biological replicates, a minimum of five technical replicates each. Error bars = +/− SD. The statistical significance was by unpaired t-test. (C) LDH release from tumor spheroids infected with WT (53%) and VNP20009 (36%) at 16 HPI. The bars represent the means of three biological replicates, a minimum of five technical replicates each. Error bars = +/− SD. Statistical significance was by one-way ANOVA with Sidak’s multiple comparison test. For all, non-significant (ns), P-value *≤ 0.05, **≤ 0.01, ***≤ 0.001, ****≤ 0.0001. (D) Representative immunofluorescence images of 3D tumor spheroids with Ki47 AlexaFluor 488 (green), Hoeschst (blue) and mCherry-tagged bacterial strains (red) at 10x and 100x objectives. Maximum intensity projections of Z-stack images were taken with an Olympus Super Resolution Spinning Disk SpinSR-10 microscope. (E) Zoomed 300% magnification of white box ROI from 100x tumor spheroids edge of both WT and VNP20009 to compare and visualize the cell morphology defect of VNP20009, outlined by dotted lines.

Representative images of VNP20009-infected tumor spheroids highlighted the reduction in intracellular bacteria compared to WT (Figure 6D), corroborating the bacterial enumeration dataset (Figure 6A, B). Furthermore, tumor spheroids infected with VNP20009 revealed distinct cell morphological defects of the strain with elongated bacterial cells (Figure 6E), confirming the strain growth defects within the tumor spheroid (see Discussion).

### Reduced CAM tumor colonization of VNP20009 compromises its anti-tumor immunity

We examined the tumor colonization ability of VNP2009 in the CAM tumor model. The effect of VNP20009 administration (∼50 CFU) on chick embryo mortality was first evaluated against the WT and PBS control groups. A significant improvement in survival rate and prolonged survival time was noted for tumor-bearing chick embryos infected with VNP20009 compared to the WT (Figure 7A). Chick embryos completely succumbed to WT infection by four days, whereas VNP-infected embryos survived for up to seven days. We next assessed the tumor-colonizing capability of VNP20009. Live *in ovo* images of mCherry-tagged bacterial strains colonizing the CAM tumor highlighted the marked reduction in fluorescence signals of VNP20009 at 1 DPI (Figure 7B) compared to the WT. The image emphasizes the strain attenuation of VNP2009 in tumor colonization, which was also apparent when we examined the intra-tumoral distribution (Supplementary Figure 5). Compared to WT, VNP20009 was remarkably reduced throughout the tumor tissue sections at all depths. This result was further confirmed by an overall reduction in the extracellular bacterial load recovered from VNP20009-infected tumors (Figure 7C). At 1 DPI, the extracellular bacterial load of VNP20009 was only ∼1% of WT. A reduction of VNP20009 tumor colonization was still evident at 3 DPI and was ∼36% of WT colonization (Figure 7C). When we compared the two strains and their ability to colonize the liver (Figure 7D), VNP2009 exhibited an early defect in dissemination (∼10% of WT). However, the attenuated strain was able to colonize chick embryo livers to a similar level as the WT by 3 DPI (Figure 7D). This was in contrast to tumor colonization (Figure 7C), where the rate of VNP2009 colonization remained weaker than the WT over time. Furthermore, a cell morphological defect was observed, as elongated chains of VNP20009 were detected in CAM tumor tissue sections at 3 DPI (Figure 7E). These chains were strikingly similar to our previous observations in 3D tumor spheroids (Figure 6E). The CAM tumor invasion rate of VNP20009 was also compromised compared to the WT invasion rate (see intracellular bacterial counts, Figure 7F).

**Figure 7:**
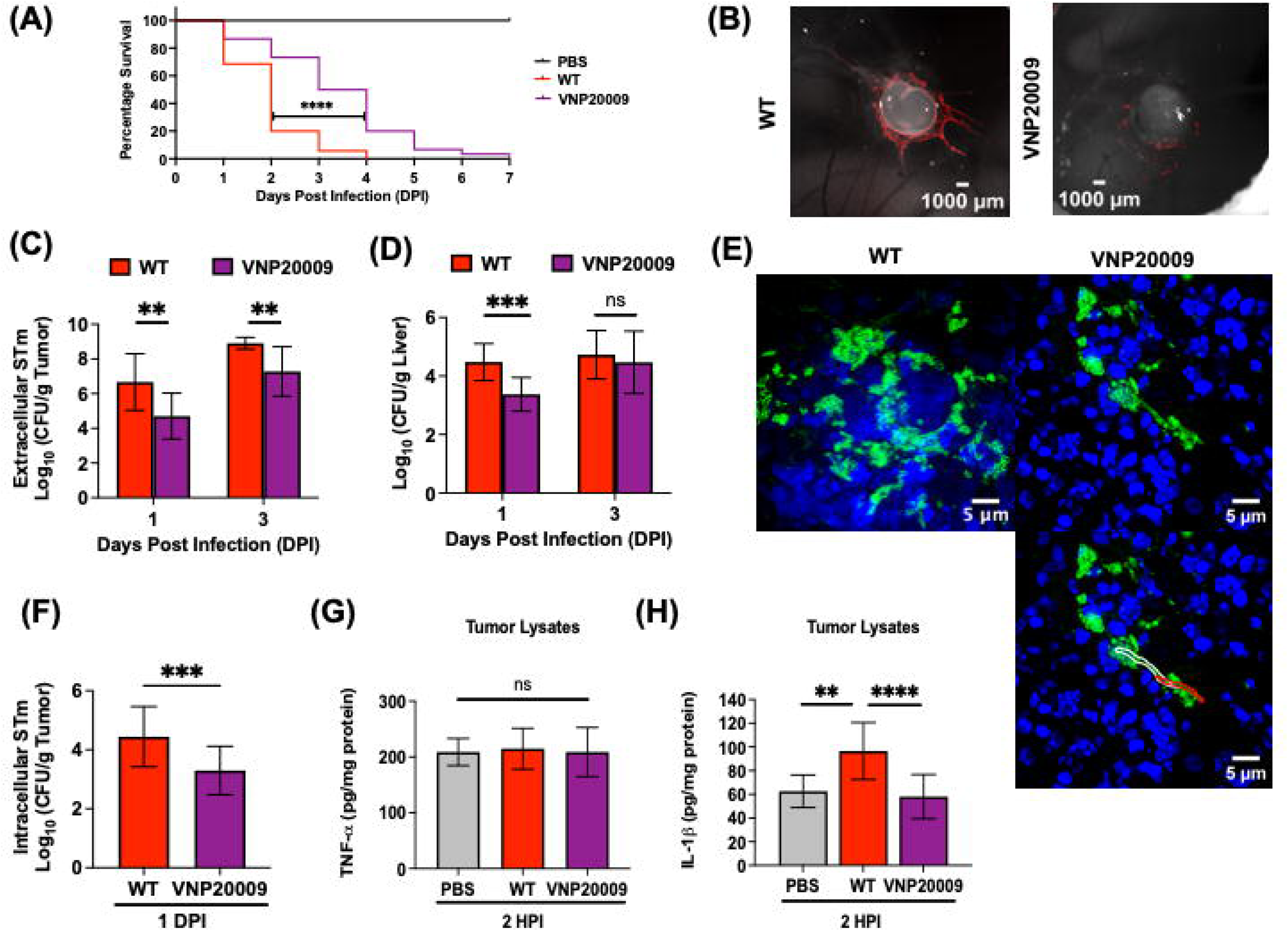
R**e**duced **tumor colonization by VNP20009 impacts its anti-tumor immunity.** (A) Tumor bearing chick embryos infected with VNP20009 (purple line) showed significant improvement in the percentage survival as well as longer survival times compared to chick embryos that were infected with WT (red line) at a dose of ∼50 CFU. For survival analysis, the Log-rank (Mantel-Cox) test was used. (B) Representative live, *in ovo* images of mCherry-tagged *Salmonella,* WT and VNP20009 colonizing CAM tumors at 1 DPI. (C-D) Extracellular *Salmonella* WT and VNP20009 CFU recovered per gram of tumor and liver at 1 and 3 DPI. The statistical significance was determined by unpaired t-test. (E) Super Resolution 320x magnified representative images of WT compared with distinct cell morphology defects of VNP20009 observed within CAM tumor tissue sections at 3 DPI. LPS (green) and Hoechst (blue). Elongated bacteria cells are outlined with white and red lines. (F) Intracellular *Salmonella* WT and VNP20009 CFU recovered per gram of tumor at 1 DPI. The statistical significance was determined by unpaired t-test. (G-H) TNF-α and IL-1β expression level in tumor lysates at 2 HPI, measured by ELISA, data were expressed as pg per mg of total protein. The results represent the mean ± SD. Statistical significance was by one-way ANOVA with Sidak’s multiple comparison test. Not significant (ns), P-value * ≤ 0.05, ** ≤ 0.01, *** ≤ 0.001, **** ≤ 0.0001. N = 2 with 5-10 tumor-bearing chick embryos per group.

Chick embryos possess only partial immunity at early developmental stages (before ED 9), but as immunocompetence develops, embryos are capable of mounting an inflammatory response with cytokine expression from host immune cells (37). Induction of pro-inflammatory cytokines such as tumor necrosis factor-alpha (TNF-α) and Interleukin-1 beta (IL-1β) following *Salmonella* infection in tumor-bearing hosts are indicative of host-mediated tumoricidal effects (38). To investigate whether host immune factors play an indirect role in driving anti-tumor immunity, CAM tumors were infected with WT, VNP20009 or PBS at ED11, and cytokine expression in tumor lysates at 2 HPI was measured. TNF-α levels were similar across uninfected and infected tumors (WT and VNP20009) at 2 HPI (Figure 7G). In contrast, IL-1β was highly expressed in WT-infected tumors, while IL-1β levels were similar between uninfected (PBS) and VNP20009-infected tumors at 2 HPI (Figure 7H). This result suggests that reduced tumor colonization by VNP20009 failed to trigger activation of host-mediated anti-tumor immunity in the CAM model.

## DISCUSSION

To the best of our knowledge, this is the first study to exploit the CAM tumor model as an *in vivo* model to study *Salmonella*-mediated anticancer therapy. Our work established that the chick embryo CAM model reproduces key features of tumor-targeting STm reported in the mouse model: (1) tumor specificity in colonization (Figure 2A), (2) intra-tumoral distribution (Figure 2B), (3) direct induction of tumor cell death (Figure 2C-G) and (4) indirect stimulation of anti-tumor immunity to some degree in the partial immunocompetent host (Figure 7H). To assess whether the CAM model recapitulated the published mouse studies, *Salmonella* key pathogenic traits were investigated for their respective role in tumor colonization and invasion.

Flagellar-dependent motility as a driver of bacterial migration, tumor colonization, and intra-tumoral distribution was not essential, as a non-motile *Salmonella* (Δ*flgK)* was not impaired for colonization in the CAM model (Figure 3). This finding agrees with published mouse data indicating that tumor colonization by *Salmonella* is a passive process (not requiring bacterial motility), where bacterial infection induces a cytokine storm and hemorrhage, facilitating the deposition of bacteria at the tumor site (23, 39). In the CAM tumor model, the passive tumor migration process of STm (via a distant inoculation route, Figure 3A, C) was most likely aided by the active movement motion of a live chick embryo just beneath the CAM layer.

In our *in vitro* 3D tumor spheroids, both SPI-1 and SPI-2 contributed to STm intracellular replication within the tumor spheroids (Figure 4). In the mouse model, there are conflicting reports as to whether SPI-1 or SPI-2 virulence genes are required for tumor invasion. Although SPI-1-mediated tumor invasion (40) and SPI-2-associated intra-tumoral replication (41) were reported to be important, in the majority of mouse tumor studies, virulence functions were not required, as bacteria were extracellular on tumor sites (23, 34, 42, 43). Consistent with these latter findings, virulence functions were not required for CAM tumor colonization and invasion (Figure 5A, B). In our experiments, *Salmonella* strains (WT, Δ*hilD,* Δ*ssrB*) were predominately extracellular on the CAM tumor (Figure 5C). The inconsistency between *in vitro* and *in ovo* data is likely attributed to the fact that 3D spheroids consist of a single cancer cell-type, which lacks the heterogeneity and vascularization of the chick CAM model to mimic the complexity of the tumor microenvironment within the host and its interaction with bacteria.

In the mouse model, an almost exclusive extracellular residence as biofilms on solid tumors was reported to be a bacterial protective mechanism against phagocytosis by host immune cells, specifically neutrophils (34). However, the CAM tumor model did not reproduce the STm biofilm phenotype, as the Δ*csgD* strain was able to colonize tumors to similar levels as WT (Figure 5D). Immunological differences in the tumor microenvironment between the two models (mouse vs chick embryos) might explain the differences in an adaptation of bacterial lifestyles on solid tumors. In contrast to a mammalian host, chick embryos have heterophils, the avian equivalent of neutrophils, but lack myeloperoxidase (37). Previously it was shown that heterophil recruitment to the inflammation site following lipopolysaccharide stimulation was detectible at ED7 (44), but fully functional heterophils within developing embryos was only described at ED18, three days before hatching (45, 46). Our CAM experimental timeline is completed by ED14 due to the low survival percentage of tumor-bearing chick embryos at an STm infectious dose of ∼50 CFU. Therefore, the lack of functional heterophils within the chick embryonic immune system in the current study timeline likely influences the non-biofilm residence of STm on CAM tumors. As with every model, the CAM tumor model has its limitations. As highlighted above, the lack of a functional host immune system and the short experimental timeline (inability to assess STm impact on tumor growth or regression) should be considered during experimental design and data interpretation.

Previous studies examining the anti-tumor efficacy of the highly attenuated STm strain VNP20009 failed to directly compare the tumor colonizing level of the strain relative to its parental WT (15, 16, 18, 47–49). Here, we used both the *in vitro* 3D tumor spheroids and CAM tumor model to provide evidence of reduced tumor colonization, invasion and replication by VNP20009 compared to WT (Figures 6AB, 7BC, F). In 3D tumor spheroids, the reduced intracellular replication of VNP20009 has consequences that limit its ability to kill tumor cells (Figure 6C). In addition, we observed cell morphology defects of VNP20009 (i.e., elongated bacteria cells) in both models during infection (Figures 6E, 7E). Although deletion of *msbB* in VNP20009 has been reported to exhibit severe growth and morphology defects under *in vitro* growth conditions in the presence of 5% CO_2_, acidic pH and high osmolarity (50, 51), our work provides additional evidence of the inherent growth sensitivity of VNP20009 in the *in vivo* tumor microenvironment (Figure 7E). We reason that this growth defect is related to cancer metabolism, which undergoes aerobic glycolysis (also known as the Warburg effect), resulting in elevated CO_2_ levels and lactic acid in the tumor microenvironment (52).

A poor human clinical outcome of VNP20009 was mainly the result of low tumor colonization that compromised its anti-tumor efficacy (20). In our CAM tumor model, we showed that IL-1β expression levels in tumor lysates of VNP20009 and PBS controls were similar (Figure 7I), suggesting that the reduced presence of VNP20009 on tumors was insufficient to elicit the anti-tumor immune response in the CAM model. These results highlighted the importance of maximizing the tumor colonizing capability of the bacterial strain as a primary factor in strengthening its anti-tumor effectiveness.

In summary, the chick embryo CAM model offers a valuable tool for initial screening and preclinical assessment of tumor-colonizing *Salmonella* strains. Its accessibility, vascularization, and relevance to tumor biology make it a suitable choice for studying bacterial interactions with tumors and assessing potential therapeutic strains for bacterial-mediated cancer therapy.

## MATERIALS AND METHODS

### 3D Spheroid infection model

A549 tumor cell spheroids were formed with the 96-well bioprinting kit (Greiner Bio-One catalog. 655841) according to the manufacturer’s guidelines. Cells were first magnetized with NanoShuttle-PL, nanoparticles overnight. Magnetized cells were counted and seeded at the same cell density of ∼50,000 cells/well. Using the 96-well magnetic drive placed at the bottom of the well, magnetized cells were printed into spheroid for 24 h. The next day, STm strains grown under SPI-1 inducing conditions were added into each well at an MOI of 100. Bacteria were added for 30 min, high gentamicin concentrations of 100 μg/ml were added for 1.5 h for invasion (2 HPI) and changed to low gentamicin of 20 μg/ml for the remainder of the infection process. We confirmed that all extracellular bacteria were eliminated post-gentamicin treatment by plating the cell culture medium. At each step of the cell media replacement and wash process, the 96-well plate was placed on the holding drive to hold the spheroids down while aspirating the solution to prevent cell loss. At both 2 HPI (invasion) and 16 HPI (replication), cell media were aspirated, washed with phosphate-buffered saline, PBS (Roche, 11666789001), and replaced with 1% Triton X-100 (Sigma, T8787-100ml) for 10 min to lyse the cells. Spheroids were homogenized by the vigorous pipetting method. Lysates were then serially diluted and plated for intracellular bacterial enumeration.

### Whole-mount Immunofluorescence staining of 3D tumor spheroids

At the end of a 16 h STm infection, tumor spheroids were fixed in 4% paraformaldehyde for ∼5 h, washed with PBS thrice, and then processed for whole-mount immunofluorescence staining. Each tumor spheroid was individually picked up using the magpen with teflon caps (Catalog no: 657850, Grenier, Bio-One), during each step of the staining process for solution replacement. Antigen retrieval was performed by transferring spheroids into a citrate buffer (Sigma, C9999-100ml) solution for 20 minutes in 99°C water bath before allowing the solution to cool to room temperature. The permeabilization step was performed with 0.3% Triton-X/PBS for 15 min followed by incubation with blocking buffer, 5% bovine serum albumin in PBS (BSA; Sigma-Aldrich, Catalog A7030) for 1 h. Spheroids were then incubated with primary antibody, recombinant anti-Ki67 (Abcam Catalog ab16667) diluted in blocking solution (1:200), overnight at 4°C in the dark. The next day, spheroids were incubated with a secondary antibody to Ki67, Alexa Fluor 488 goat anti-rabbit IgG H&L (Abcam Catalog ab150081), and Hoechst (blue, Invitrogen, H3570) for 1 h. Whole spheroids were then mounted onto the slide with a prolong gold antifade mountant (Invitrogen Catalog: P36934), coverslip sealed with nail polish and dried overnight in the dark. Whole-mount spheroids were imaged with Olympus SpinSR-10 spinning disk confocal at 10x air objective (NA 0.40), and 100x oil objective (NA 1.5). Acquired microscopy images were processed with Fiji software (25).

### Embryonated chicken CAM tumor model

Specific pathogen-free (SPF), premium fertilized chicken eggs were purchased from AVS bio and set horizontally in a Rcom Max 50 incubator at 37°C and 60% humidity. On ED3, CAM of the chick embryo was dropped by inserting a thin needle at the apex of the egg and removing ∼4 ml of albumin with a syringe and 21G needle. A small window (∼1 cm^2^) was made on top of the eggshell and resealed with 3M Tegaderm TM transparent film dressing. On ED7, A549 cell pellets (∼2 x 10^6^ cells/egg) were mixed with 50 µl Matrigel (Corning 354234 or 354263) and were grafted into the center of a sterilized silicone ring placed on top of the CAM layer. Eggshell windows were then re-sealed with Tegaderm and put back into the incubator to allow tumor growth for four days. On ED11, the silicone ring was first carefully removed from the CAM without disrupting the established tumor before the application of the bacterial inoculum directly onto the tumor with a pipette. Bacterial inoculums were plated each time to confirm that a similar dose was applied for all strains. For the distant inoculation route experiment, bacteria were deposited onto a sterile cut-out filter paper, placed a short distance away from the tumor (∼0.5 cm). At different time points, tumors and livers were harvested for enumeration of viable bacteria by homogenization, serial dilution, and plating methods. All chick embryo experiments were repeated twice (N = 2) with 5-10 tumor bearing embryos in each treated group.

### TUNEL assay and apoptosis quantification with Fiji

Detection of apoptotic cells on tumor tissue sections was performed with Click-iT Plus TUNEL assay (Invitrogen, Thermo Fisher Scientific Lot 2559160) according to the manufacturer’s guidelines. Immunofluorescence images were acquired with an Olympus Super Res Spinning Disk, SpinSR-10 microscope at 100x oil objective (NA 1.5). Quantification of TUNEL-positive nuclei as well as DAPI nuclei was performed with Fiji (25) following the optimized and unbiased image processing protocol. Briefly, images were first split into separate color channels and compressed with z-axis maximum intensity projection. Background noise reduction was performed by subtracting background values from each channel. Images were then converted to greyscale with 8-bit. The Binarization thresholding function available in Fiji (25) was utilized and thresholding algorithms that capture the best outline of stained cells for each channel were selected (53). Moments and Li auto-thresholding algorithms were opted for highlighting TUNEL-positive nuclei and DAPI-positive cells, respectively. Lastly, the analyze particle function was applied for the count read-outs. The percentage of TUNEL-stained cells was calculated against the total cell numbers (DAPI-positive cells) within the same region.

### Statistical Analysis

All statistical analysis were performed with GraphPad Prism 10.2.1.

## Supporting information

Supplemental Figures and Method Details

## Acknowledgements

We are extremely grateful for the thoughtful comments provided by our reviewers. We thank Dr. Ruby Yun-Ju Huang from National University of Singapore for guidance and expertise on the chick chorioallantoic membrane (CAM) tumor model. This work was supported by Cancer prevention and Research Institute of Texas (CPRIT) Grant RP2000650 awarded to LJ.K.

## Abbreviations

STm: *Salmonella* Typhimurium
MOI: multiplicity of infection
ED: embryonic development day
DPI: days post-infection
HPI: hours post-infection
CAM: chorioallantoic membrane
LDH: Lactate Dehydrogenase
H&E: Hematoxylin and Eosin
WT: wildtype

