## Supplemental Figures and Method Details for "Evaluation of Tumor-colonizing *Salmonella* Strains using the Chick Chorioallantoic Membrane Model"

| **Strain or plasmid** | **Description** | **Source or reference** |
| --- | --- | --- |
| Wildtype | *Salmonella enterica* serovar Typhimurium 14028s, pFPV::mCherry | Lab strain collection |
| Δ*flgK* | *flgK::cm,* pFPV::mCherry | Lab strain collection |
| Δ*hilD* | *hilD* deletion, pFPV::mCherry | Lab strain collection |
| Δ*ssrB* | *ssrB::kan,* pFPV::mCherry | Desai et al. (2016) |
| Δ*csgD* | *csgD::chl,* pFPV::mCherry | Desai et al. (2016) |
| VNP20009 | *Salmonella enterica* subsp. *enterica* strain YS1646, pFPV::mCherry | ATCC BAA3199 |
| pFPV::mCherry plasmid | mCherry cloned between XbaI and SphI sites in pFPV | Addgene plasmid # 20956  http://n2t.net/addgene:20956; RRID:Addgene_20956 |
| pWSK29-P*prgH*- mCherry | plasmid pWSK29 containing the *prgH* promoter fused before mCherry | Lab strain collection |

**Supplementary Table 1: List of bacterial strains and plasmid used in the study.**

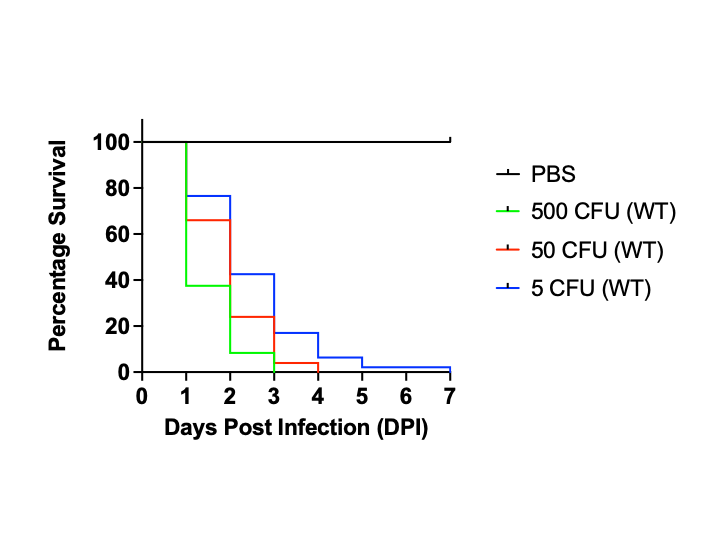

**Supplementary Figure 1:** Optimal infectious dose of STm for the CAM tumor model was approximately 50 CFU. Dose response survival analysis was performed by infecting the tumor bearing chick embryos with different infectious doses of STm (~500 CFU, ~50 CFU, ~5 CFU) or PBS alone. Mortality of chick embryos were monitored and recorded daily following STm infection. Death of the chick embryo was identified through a small window opening on the top of an eggshell (covered with transparent Tegaderm) and was characterized by the loss of blood vessels and non-movement of the chick inside the egg.

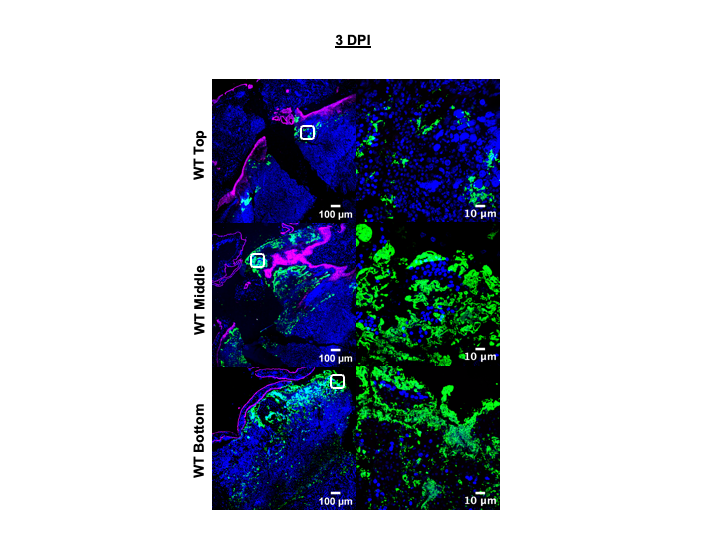

**Supplementary Figure 2:** Representative images of STm WT LPS (green), E-cadherin (magenta), and Hoechst (blue) staining in tumor sections from top, middle, and bottom depth levels of whole tumors harvested at 3 DPI. Scale bar = 100 µm (10x objective) and 10 µm (100x objective). The second panel is the zoomed image of the white box at 100x objective.

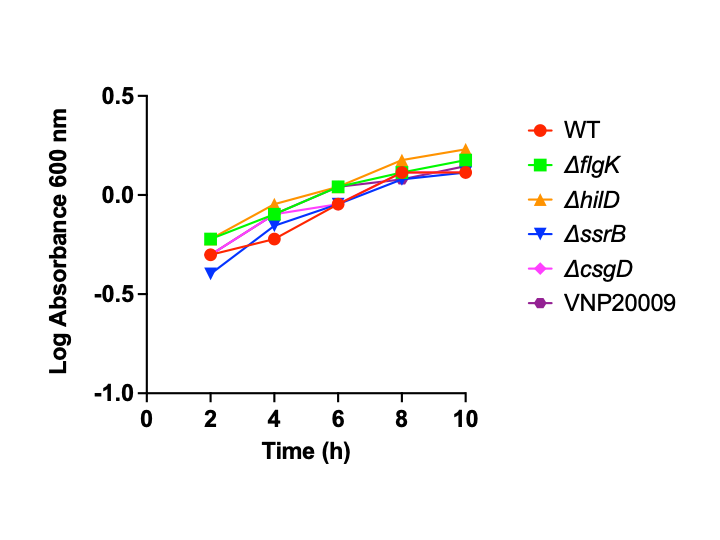
**Supplementary Figure 3:** VNP20009 was not defective for growth under lab growth conditions. STm strains deleted in flgK, hilD, ssrB, csgD, VNP20009, and the wild type 14028s were grown for 10 h in Luria-Bertani broth at 37°C, 250 rpm and the absorbance at 600 nm was measured every 2 h.

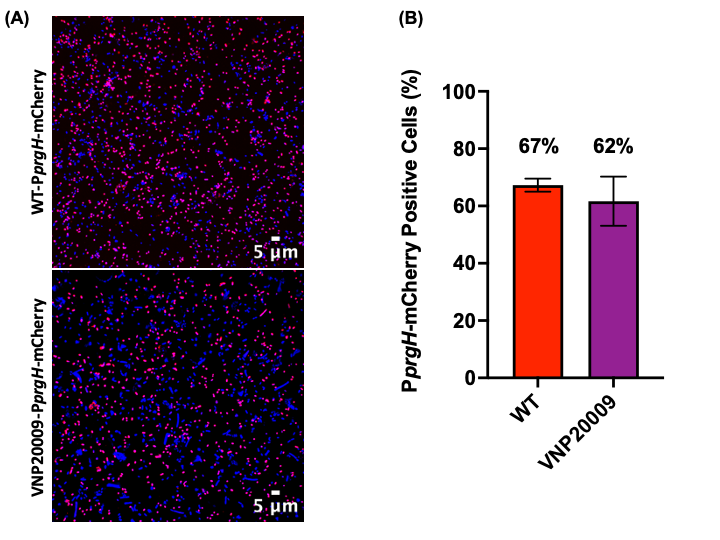

**Supplementary Figure 4:** SPI-1 expression (using a P*prgH*-mCherry reporter) in STm WT and VNP20009 strains, grown under SPI-1 inducing condition. **(A)** Representative images of STm WT- and VNP20009-P*prgH*-mCherry stained with Hoechst (blue). (B) Quantification comparison of P*prgH*-mCherry positive cells between WT (67%) and VNP20009 (62%), of the total population (Hoechst, blue). The results represent the mean ± SD.

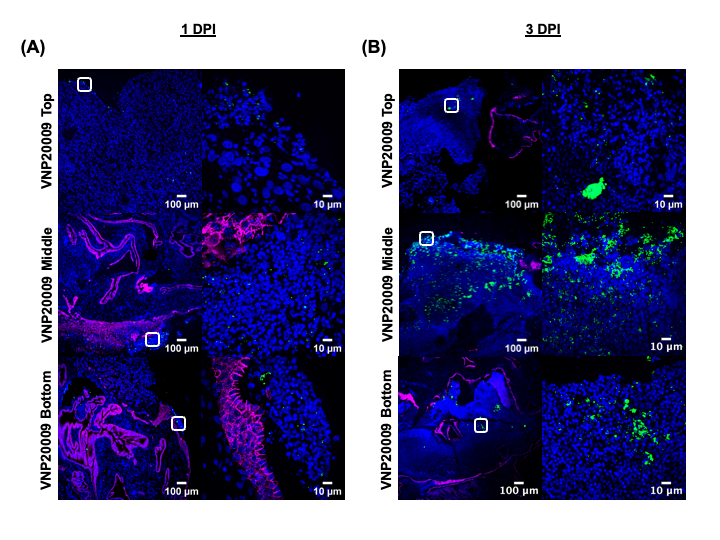

**Supplementary Figure 5:** Representative images of STm VNP20009 LPS (green), E-cadherin (magenta), and Hoechst (blue) staining in tumor sections from top, middle, and bottom depth levels of whole tumors harvested at (A) 1 DPI and (B) 3 DPI. Scale bar = 100 µm (10x objective) and 10 µm (100x objective). The second panel is the zoomed image of the white box at 100x objective.

**METHOD DETAILS**

**Cell culture and bacterial strains growth conditions**

A549 human lung carcinoma cells (ATCC CCL-185) were purchased from the American Type Culture Collection (ATCC). The cell line was cultured in DMEM (Gibco 11965-092) supplemented with 10% heat-inactivated fetal bovine serum (GenDEPOT, F0900-050) and 1% penicillin/streptomycin (GenDEPOT, CA005-010), at 37°C with 5% CO_2_. The bacterial strains and plasmid used in this study are listed in Supplementary Table 1. Bacterial strains (transformed with reporter plasmid) were grown in Luria-Bertani (LB) medium in presence of 100 μg/ml ampicillin (Sigma, A0166) at 37°C at 225 rpm overnight. Sub-cultures were grown to the late exponential phase to induce SPI-1 associated genes and diluted to the desired dosage for inoculum preparation in both 3D tumor spheroids and CAM tumor models.

**Lactate Dehydrogenase Assay**

The cell culture supernatants from A549 tumor spheroids infected with STm strains were collected at 16 h post-infection times and a Lactate Dehydrogenase (LDH) assay was performed with CytoTox96 non-radioactive cytotoxicity assay according to the manufacturer’s guidelines (Promega, Catalog no: G1780). Maximum releasable LDH activity (100%) in the cells was achieved by treating the uninfected spheroids with lysis buffer for 45 min at room temperature and the maximum LDH release control measurement was used to calculate the percentage of cytotoxicity in STm-infected cells. The colorimetric absorbance of the samples was measured with a plate reader at 490 nm (Bio-rad iMark microplate absorbance reader #1681130).

**CAM tumor dissociation for tumor cell viability**

Whole tumors were harvested and placed in a sterile petri dish where they were minced with scissors in DPBS++ solution (Corning 21-030-CV). Minced tissue fragments with the solution were transferred into 15 ml centrifuge tubes. DPBS++ solution was removed by centrifugation and replaced with 2 ml (1mg/ml) of collagenase (Millipore Sigma SCR103). Tubes were turned over several times and incubated at 37°C for 3-5 h. At the end of incubation, 5 ml of DMEM media (without antibiotic) were added to each tube and complete tissue cell dissociation ensured by pipetting from 20 to 40 times. Tissue fragments with media solution were filtered through a 40 µm nylon mesh cell strainer (Falcon 352340). Filtered cell suspensions were then centrifuged at 500 x g for 5 min. Supernatants were removed and the dissociated cell pellet was resuspended with fresh DMEM. The total viable tumor cell count was obtained with the trypan blue staining method.

***Ex vivo* gentamicin protection assay**

To evaluate the percentage of intracellular bacteria in the tumor tissue, an *ex vivo* gentamicin protection assay was performed. The whole tumor tissue was homogenized in 1 ml of cold PBS. A 0.5 ml aliquot of this suspension was then serially diluted and plated to determine the extracellular bacterial load in the tumor. The remaining 0.5 ml aliquot was treated with 200 µg/ml of gentamicin for 90 min at 37°C to kill extracellular bacteria. Tissue suspensions were then centrifuged at 5,000 rpm for 10 min, washed twice with PBS, and resuspended in 1% Triton X-100 PBS for 15 min at 37°C to lyse the cells. The lysate was then serially diluted and plated for the intracellular bacteria enumeration.

**Histology and immunofluorescence staining**

Tumors harvested at 1 and 3 DPI were fixed in 10% formalin (Sigma, H7501128) before samples were sent to the UTMB histopathology core facilities for paraffin embedding, tissue sections as well as Hemolysin and eosin (H&E) staining. H&E-stained tumor sections were sent to the Research Histology Core Laboratory (RHCL) at MD Anderson for external histopathologist assessment. The sample thickness was 5 µm at each depth: top, middle, and bottom cuts of the tumor samples. Tissues were deparaffinized and rehydrated in a series of xylene (Sigma, 534056-500ML) and ethanol (Fisher Scientific, BP2818500) washes. Deparaffinized and rehydrated tumor sections were boiled in antigen retrieval solution, citrate buffer for 35 min at 99°C water bath before allowing the solution to cool to room temperature. Permeabilization was performed with 0.3% Triton-X/PBS for 15 min followed by incubation with blocking buffer, 5% bovine serum albumin in PBS for 1 hr. Tumor sections were then stained and prepared for immunofluorescence with LPS (Abcam, ab35156) 488 (green) for Salmonella, E-cadherin (BD Biosciences, 610182) 647 (far-red), and Hoechst (blue) for the nucleus. Images were acquired with an Olympus Super Res Spinning Disk, SpinSR-10 microscope using 100x oil objective (NA 1.5) and super resolution imaging with optical zoom, 320x of 100x region of interest (ROI) for VNP20009 elongated cells observation.

**ELISA**

Tumor tissues were collected at 2 h post-infection (HPI) and homogenized in Radioimmunoprecipitation assay (RIPA) buffer (Cell Signaling Technology, 9806S) containing protease inhibitor cocktail (Roche, 11836153001). Protein lysates were collected from supernatant after a round of centrifugation at 13,000 x g for 20 min, 4°C and were stored immediately at -80°C until analysis. Total protein concentration from tumor lysates was measured with Pierce BCA assay (ThermoScientific 23225) following the manufacturer’s instructions. Tumor lysates were normalized for total protein content and analyzed by chicken-specific enzymatic-linked immunosorbent assay (ELISA) kits for expression levels of TNF-alpha (MyBioSource MBS:260419) and IL-1beta (MyBioSource MBS:2024496) according to the manufacturer’s guidelines.
